# E3 ubiquitin ligase MARCHF5 controls BAK apoptotic activity independently of BH3-only proteins

**DOI:** 10.1101/2022.01.04.474880

**Authors:** Shuai A. Huang, Hui San Chin, Boris Reljic, Tirta M. Djajawi, Iris K.L. Tan, David A. Stroud, David C.S. Huang, Mark F. van Delft, Grant Dewson

## Abstract

Intrinsic apoptosis is principally governed by the BCL-2 family of proteins, but some non-BCL-2 proteins are also critical to control this process. To identify novel apoptosis regulators, we performed a genome-wide CRISPR-Cas9 library screen, and it identified the mitochondrial E3 ubiquitin ligase MARCHF5/MITOL/RNF153 as an important regulator of BAK apoptotic function. Deleting MARCHF5 in diverse cell lines dependent on BAK conferred profound resistance to BH3-mimetic drugs. The loss of MARCHF5 or its E3 ubiquitin ligase activity surprisingly drove BAK to adopt an activated conformation, with resistance to BH3-mimetics afforded by the formation of inhibitory complexes with pro-survival proteins MCL-1 and BCL-XL. Importantly, these changes to BAK conformation and pro-survival association occurred independently of BH3-only proteins and influence on pro-survival proteins. This study identifies a new mechanism by which MARCHF5 regulates apoptotic cell death and provides new insight into how cancer cells respond to BH3-mimetic drugs. These data also highlight the emerging role of ubiquitin signalling in apoptosis that may be exploited therapeutically.

## INTRODUCTION

Intrinsic apoptosis plays a pivotal role in tissue homeostasis, development and immunity. Dysregulation of apoptosis contributes to a variety of human pathologies such as neurodegenerative disorders ^1^, autoimmune diseases ^2^ and cancers ^3^. Therefore, targeting apoptosis is emerging as an attractive strategy to either block cell death in neurodegenerative disease or to induce cell death to treat cancer. In vertebrates, intrinsic apoptosis is principally regulated by BCL-2 proteins, which comprise three subgroups: the pro-survival proteins, the pro-apoptotic effector proteins and the initiator BH3-only proteins. These proteins control cellular fate through protein-protein interactions ^4,5,6^, with some associations promoting death and others favouring survival.

Although apoptosis is mainly controlled by BCL-2 proteins, some non-BCL-2 proteins also serve as important regulators ^7,8,9^. These include voltage-dependent anion channel 2 (VDAC2), which supports the integration of the cell death effector proteins BAK and BAX into the mitochondrial outer membrane (MOM) ^10,11,12,13^.

In healthy cells, BAK and BAX molecules that reside on the MOM associate with VDAC2 ^13^. However, these interactions with VDAC2 have seemingly opposing effects on BAX and BAK apoptotic function. Whilst deleting *Vdac2* inhibits BAX apoptotic activity by impairing BAX localisation to the MOM ^14^ it elevates BAK apoptotic activity ^15^, even though BAK levels at mitochondria are markedly reduced in *Vdac2*-deficient cells ^11, 13, 14^. These findings emphasize that although BAX and BAK are typically regarded as redundant apoptosis effector proteins, they are regulated distinctly from one another.

To understand why BAK exhibits heightened activity in the absence of VDAC2, we performed a CRISPR-Cas9 whole genome library screen that identified the mitochondrial E3 ubiquitin ligase MARCHF5 as a novel regulator of BAK-dependent apoptosis. Loss of MARCHF5 rendered BAX-deficient cells highly resistant to prolonged treatment with BH3 mimetics. Abolishing MARCHF5 E3 ubiquitin ligase activity was sufficient for this phenotype, which involved BAK adopting an active conformation and forming inhibitory Mode 2 complexes with the pro-survival proteins MCL-1 and BCL-XL. Surprisingly, these changes to BAK conformation and association did not require BH3-only proteins.

## METHODS and MATERIALS

### Cell Culture and Stable Cell Lines

SV40-transformed Mouse Embryonic Fibroblasts (MEFs) were cultured in Dulbecco’s Modified Eagle’s Medium (DMEM) supplemented with 10% (v/v) Fetal Bovine Serum (FBS), 50 μM 2-Mercaptoethanol (2-ME) and 100 μM asparagine. HeLa cells (ATCC#CCL-2) and Human Embryonic Kidney 293T (HEK293T) cells (ATCC #CRK-3216) were cultured in DMEM supplemented with 10% (v/v) FCS. HCT-116 (gift from Luo’s Laboratory ^16^) and KMS-12-PE (DSMZ #ACC606) cells were cultured in RPMI1640 supplemented with 10% (v/v) FCS. Cells were cultured in humidified incubators maintained at 37 °C and 10% CO_2_ for MEFs, HeLa and 293T cells or 5% CO_2_ for KMS-12-PE cells. Cells were routinely (every two months) screened for Mycoplasma contamination by MycoAlert Kit (Cat#LT07218, Lonza) as the manufacturer’s introductions. Only Mycoplasma negative cells were analysed.

### Retroviral and lentiviral infection

3xMyc (3xMEQKLISEEDLNE)-tagged wild-type MARCHF5 or MARCHF5C65/68S were synthesised as gene fragments by Integrated DNA Technologies (IDT) and subcloned into pMSCV-IRES-puro (pMIP) vector ^17^ or pMSCV-IRES-hygro (pMIH) vector. HEK293T cells were used as packaging cells. The retroviral or lentiviral constructs were first introduced into packaging cells by FuGENE6 (Cat#E2691, Promega) transfection according to the manufacturer’s instructions. Viral supernatants were filtered and used to infect MEF, HeLa and KMS-12-PE cells by spin-infection (2500 rpm centrifugation at 25 °C for 1 hour) in the presence of 4 µg/mL polybrene (Cat#S2667, Sigma-Aldrich). Stably transduced cells were selected either by antibiotic selection (2 mg/mL hygromycin (Cat# 10687010, Thermo Fisher) or 2 µg/mL puromycin (Cat#P9620, Sigma-Aldrich)) or expression of fluorescent protein markers (GFP, BFP or CFP) using a BD FACSAria Fusion cell sorter.

### Generating knockout cell lines by CRISPR/Cas9 gene editing

Gene-deletions were engineered using CRISPR-Cas9 gene editing technology either by transient transfection or lentiviral transduction. The sgRNA sequences used for targeting specific genes (Table S1) were taken from available whole genome libraries Gecko ^18^ or Brunello ^19^. Oligonucleotides encoding these sgRNA sequences were ordered from IDT and annealed as previously described ^20^. The annealed oligos were cloned into pSpCas9(BB)-2A-GFP/PX458 (for transient co-expression of sgRNA and Cas9) ^21^, pKLV-U6RNA(BbsI)-PGKpuro2ABFP (for constitutive lentiviral sgRNA expression) ^22^, or FgH1tUTG-CFP or FgH1tUTG-GFP vectors (for inducible lentiviral sgRNA expression) ^23^. FUCas9Cherry lentiviral vector was used to generate stable Cas9 expressing HeLa and KMS-12-PE cells ^23^. sgRNA were expressed for 48h before cells were sorted into single cells based on fluorescent marker protein expression using a BD FACSAria Fusion cell sorter. Individual clones were expanded, and successful knock-out clones were identified by sequencing PCR amplicons encompassing the sgRNA target sites. The primers for amplifying the sgRNA target sites are listed in Table S2.

### Immunoblotting and immunoprecipitation

Cell pellets were lysed in ONYX lysis buffer (20 mM Tris-pH 7.4, 135 mM NaCl, 1.5 mM MgCl_2_, 1 mM EGTA, 10% (v/v) glycerol, supplemented with 1% (v/v) Triton X-100 or 1% digitonin, 0.5 μg/ml complete protease inhibitor (Sigma-Aldrich)) for 30 min on ice. Following lysis, samples were centrifuged at 13,000 rpm at 4 °C for 10 minutes and supernatants were collected. Protein concentration was determined using the Bradford protein assay. For immunoprecipitation, bait proteins were captured with antibodies to BAK (Clone# G317-2, BD Pharmingen), MCL-1 (Clone#19C4-15, Walter and Eliza Hall Institute) or BCL-XL (Clone#E18, Abcam) and Protein G Sepharose beads. 25 μg of whole cell lysate was loaded per well and resolved by SDS-PAGE. Gels were transferred onto nitrocellulose membrane (Invitrogen) using an iBlot Gel Transfer Device (Invitrogen). Non-specific binding was blocked with 5% w/v non-fat milk in TBS-T (20 mM Tris-HCL pH 7.6, 137 mM NaCl, 0.1% Tween-20) for 1 hour at room temperature before immunoblotting with primary antibodies overnight at 4 °C. The membranes were washed with TBS-T three times, followed by incubating with appropriate HRP-conjugated secondary antibodies for 1 hour in room temperature. The membranes were washed with TBS-Tween (0.01% v/v) prior to detection using enhanced chemiluminescent (ECL) reagent (Millipore) and imaged using a ChemiDoc Imaging System (BioRad). Monoclonal antibodies to MCL-1 (Clone#19C4-15), BAK (Clone# 7D10), BCL-2 (Clone# BCL-2-100), BCL-W, BIM (Clone#3C5), BID (Clone#2D1-3), MYC (Clone#7E10), VDAC2 (gift of M. Ryan) and HSP70 (Clone#N6, gift of W. Welch and R. Anderson) were produced with the WEHI antibody facility. The following antibodies were purchased from commercial sources: BAK (Cat# B5897, Sigma-Aldrich), BCL-XL (Clone#44/BCL-X, BD Bioscience), BCL-XL (Clone#E18, Abcam), BCL-2 (Clone#7/BCL-2, BD Bioscience), MCL-1 (Cat#600-401-394, Rockland), NOXA (Clone#114C307.1, Novus Biologicals), BIM (Cat#ADIAAP330, Stressgen), BAD (Cat#ADIAAP020, Stressgen), PUMA (Cat#3043, ProSci), HA (Clone#3F10, Roche Applied Science), Cytochrome c (Clone#7H8.2C12, BD Pharmingen), Cytochrome c (Clone#6H2.B4, BD Pharmingen) and FAR1 (NBP1-89847, Novus Biologicals). Secondary antibodies: anti-rabbit IgG (Cat#403005), anti-mouse IgG (Cat#103005), anti-rat IgG (Cat#303005) were obtained from Southern Biotech.

### Cell viability assays

Cells were cultured with the indicated concentrations of BH3 mimetics (ABT-737 (Cat#A-1002, Active Biochem), A-1331852 (WEHI Chemical Biology Division) and S-63845 (Cat#A-6044, Active Biochem)). After 24h, both viable and dead cells were harvested and re-suspended in KDS-BSS buffer supplemented with 2 µg/ml propidium iodide (PI) (Sigma Aldrich). Cell viability (PI negative) was assessed using an LSR-II flow cytometer (BD Biosciences).

### Clonony formation assay

Cells were seeded in 6-well plate at a density of 1,000 cells per well and cultured with titrated concentrations of BH3 mimetics for 5 days. After treatment, cells were stained with 0.5% crystal violet (Cat#C0775, Sigma-Aldrich) for 20 min at room temperature and photographed using a ChemiDoc Imaging System (BioRad).

### BAK activation, cytochrome *c* release and caspase-3 activation by Flow Cytometry

Cells were harvested following 0.5 or 2 hours of culture with BH3 mimetic drugs. 20 µM caspase inhibitor QVD-OPh (Cat#OPH109, MP Biomedicals) was included when assays were performed for BAK activation or cytochrome *c* release, but not when caspase-3 activation was measured. Harvested cells were washed once with ice-cold PBS.

To assess BAK activation and caspase-3 activation, cell pellets were fixed and then permeabilized using the eBioscience cell fixation and permeabilization kit (Cat#88882400, Thermo Fischer) according to the manufacturer’s instructions. Fixed cells were incubated with either conformation specific BAK antibody (Clone# G317-2, 1:100) (Cat# 556382, BD Pharmingen) followed by phycoerythrin (PE)-conjugated anti-mouse antibody (1:200) (Cat#1031309, SouthernBiotech) or anti-caspase-3-phycoerythrin antibody (Clone#C92-605, 1:25) (Cat#550821, BD Pharmingen).

Cytochrome *c* release was assayed as described ^24^. Briefly, cells were permeabilized by the 0.025% w/v digitonin in permeabilization buffer (20 mM 4-(2-Hydroxyethyl)piperazine_1-ethanesulfonic acid (HEPES) pH 7.5, 100 mM sucrose, 2.5 mM MgCl_2_, 100 mM KCL) for 10 mins on ice before fixation. Fixed cells were incubated with anti-cytochrome c-APC antibody (Clone#REA702, 1:50) (Cat#130111180, Miltenyi Biotec). Samples were analyzed on an LSR-II flow cytometer (BD Biosciences).

### Mass Spectrometric analysis

MEF cells were harvested, and mitochondria were isolated and enriched using a Mitochondria isolation kit as the manufacturer’s introductions (Cat#130096946, Miltenyi Biotec). HeLa cells were harvested and permeabilized by 0.025% w/v digitonin (Biosynth) in permeabilization buffer for 10 mins on ice, and then mitochondria were isolated and enriched using gradient centrifugation ^25^. As previously described ^26^, membrane fractions were normalized for protein content using the Pierce BCA Protein Assay Kit (Cat#23227, ThermoFisher Scientific), subjected to trypsinization, and peptides were labelled with Tandem Mass Tags (Cat#TMT; 90061, ThermoFisher Scientific).

For TMT labelling, protein pellets were solubilized and digested into tryptic peptides for mass spectrometry using the iST-NHS kit (PreOmics GmbH, Planegg, Germany) as per manufacturer instructions. Peptides were labelled with 6plex Tandem Mass Tags (TMT) (Thermo Fisher Scientific) in 8:1 label:protein ratio as per manufacturer instructions. Pooled samples were fractionated using the Pierce High pH Reversed-Phase Peptide Fractionation Kit (Thermo Fisher Scientific) as per manufacturer’s instructions. Individual fractions were dried using a CentriVap Benchtop Vacuum Concentrator (Labconco) and reconstituted in 2% (v/v) acetonitrile (ACN) and 0.1% (v/v) trifluoroacetic acid (TFA). Liquid chromatography (LC) coupled MS/MS was carried out on an Orbitrap Q Exactive Plus mass spectrometer (Thermo Fisher Scientific) with a nanoESI interface in conjunction with an Ultimate 3000 RSLC nanoHPLC (Dionex Ultimate 3000). The LC system was equipped with an Acclaim Pepmap nano-trap column (Dionex-C18, 100 Å, 75 μm × 2 cm) and an Acclaim Pepmap RSLC analytical column (Dionex-C18, 100 Å, 75 μm × 50 cm). The mass spectrometer was operated in a centroid data spectrum mode. The tryptic peptides were injected to the trap column at an isocratic flow of 5 μl/min of 2% (v/v) CH3CN containing 0.1% (v/v) formic acid for 5 min applied before the trap column was switched in-line with the analytical column. The eluents were 5% DMSO in 0.1% v/v formic acid (solvent A) and 5% DMSO in 100% v/v CH3CN and 0.1% v/v formic acid (solvent B). The flow gradient was (1) 0–6min at 3% B, (2) 6–95 min, 3–22% B (3) 95– 105 min 22–40% B (4) 105–110 min, 40–80% B (5) 110–115 min, 80–80% B (6) 115–125 min, 80–3% and equilibrated at 3% B for 5 min before the next sample injection. For TMT and LFQ analysis, in brief data were collected in positive mode using Data Dependent Acquisition using m/z 375–1450 as MS scan range, HCD for MS/MS of the 12 most intense ions with charge ≥ 2. Other instrument parameters were: MS1 scan at 70,000 resolution (at 200 m/z), MS maximum injection time 50 ms, AGC target 3E6, normalized collision energy was at 27% energy, isolation window of 1.2 Da, MS/MS resolution 17,500, MS/MS AGC target of 5E4, MS/MS maximum injection time 50 ms, minimum intensity was set at 2E3 and dynamic exclusion was set to 30 s. Raw files were processed using the MaxQuant platform (version 1.6.5.0) and searched against UniProt human or mouse database using default settings for a TMT 6plex or LFQ experiment. In brief: oxidation of methionine and N-terminal acetylation were specified as variable modifications. Trypsin/P cleavage specificity (cleaves after lysine or arginine, even when proline is present) was used with a maximum of 2 missed cleavages. Carbamidomethylation of cysteine was set as a fixed modification. A search tolerance of 4.5 ppm was used for MS1 and 20 ppm for MS2 matching. False discovery rates (FDR) were determined through the target-decoy approach set to 1% for both peptides and proteins.

Log_2_-transformed TMT reporter intensity corrected values were grouped and fit to a moderated t-test to identify significant differences in protein abundance between samples (limma Bioconductor package - version 3.46.0). For the MEF data, as these data were derived from independent experiments analyzed on separate mass spectrometry runs, the Log2-transformed reporter intensities were normalized and batch corrected prior to fitting the linear model with limma. For the HeLa data, the label free samples were run through the mass spectrometer in a single experiment, so batch correction was not necessary, with the LFQ intensity data otherwise treated similarly to TMT reporter intensity data described above. Mass spectrometry data has been deposited to the ProteomeXchange Consortium ^27^ via the PRIDE partner repository ^28^ under accession number PXD030246.

### CRISPR/Cas9 library screens

Cells were first engineered to stably express Cas9 by transduction with FUCas9Cherry lentivirus ^23^. mCherry^high^ cells were sorted as population using a BD FACSAria Fusion cell sorter. The Cas9-expressing cells were seeded in 10 cm^2^ dishes at a density of 1 x10^6^ one day prior to transduction with lentiviral sgRNA libraries. A genome-wide library was used for the MEF screen in Figure 1C ^22^; and a custom library was used for the HeLa screen in Figure 6C&D (synthesized by Synbio Technologies), which included 3326 sgRNAs (Table S5) targeting 514 human protein-coding genes (258 potential MARCHF5 substrates identified by our mass spectrometry experiments (Figures x, y) along with other previously described MARCHF5 substrates, known mitochondrial and peroxisomal proteins, and BCL-2 family members). Transduced cells were cultured with 2μg/ml puromycin for 48h to enrich sgRNA-expressing cells. Each transduced pool of cells was then split into two groups: one was treated with DMSO, and the other was subjected to a lethal dose of BH3 mimetics (10 µM ABT-737 or 500 nM A1331852 for MEFs and 2 µM S63845 + 1 µM A1131852 for HeLa cells) for 48h. Surviving cells were expanded for 5 days, their DNA was extracted, and sgRNA sequences were amplified by PCR and sequenced as described ^22^. For the MEF experiments, the screen was performed with two independently derived cell lines, each transduced 5-6 times with library sgRNAs, then divided and selected with either ABT-737 or A1331852 in parallel, making >20 biological replicate samples. Likewise, the Hela screen was performed with triplicate viral transductions in two independent cell clones, for a total of six replicate samples. Raw counts of each sgRNA were normalized to counts per million (CPM) reads and Log_2_ transformed following the addition of a 1 CPM offset to each sgRNA. The log_2_ transformed values were fit to a linear model and a moderated t-test was performed to identify sgRNA whose relative abundance differed significantly between the BH3 mimetic and DMSO treated samples (and also between Cas9 expressing and non-expressing cells in HeLa screen) (limma Bioconductor package - version 3.46.0). The sgRNAs were then ranked in order of their t-statistic. An iterative hypergeometric scoring algorithm ^29, 30^ was applied to identify genes whose sgRNA were enriched among the most significantly depleted sgRNA in the library.

**Fig. 1.**
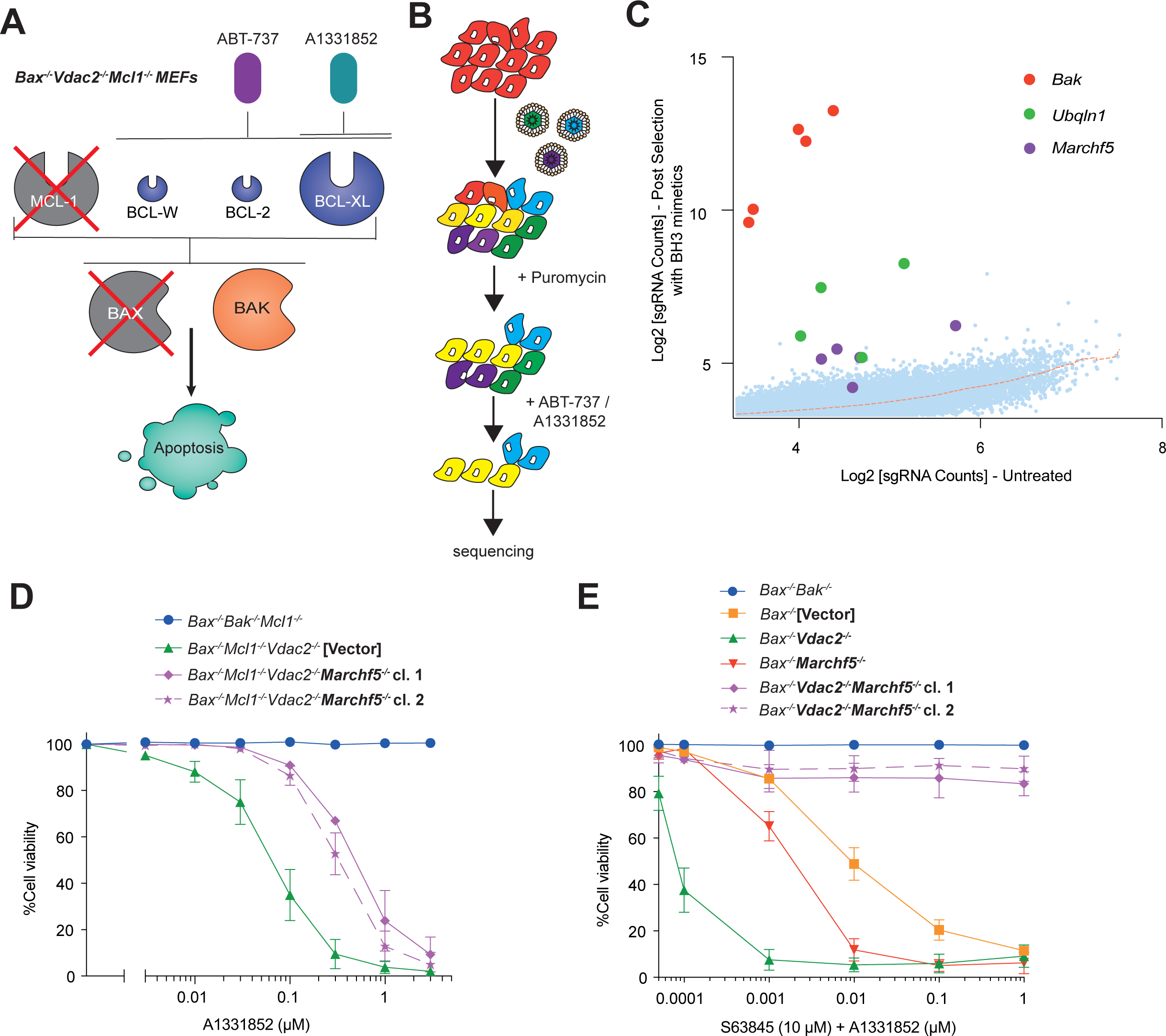
Genome-wide CRISPR/Cas9 library screen identifies MARCHF5 as a regulator of BAK-dependent apoptosis. (**A**) *Bax^-/-^Mcl-1^-/-^Vdac2^-/-^* MEFs were used for the CRISPR/Cas9 library screen. On this genetic background, apoptosis is driven exclusively by the effector protein BAK (because BAX is absent) and BAK is controlled primarily by the pro-survival protein BCL-XL (due to the absence of MCL1 - its other major regulator ^5^). Apoptosis can be provoked in these cells with BH3 mimetics that target BCL-XL, such as A1331852, which selectively inhibits BCL-XL or ABT-737, which also inhibits BCL-W and BCL-2. (**B**) Outline of the genome-wide CRISPR/Cas9 library screen. (**C**) sgRNAs targeting *Bak, Ubqln1* and *Marchf5* were significantly enriched following selection with ABT-737. *Bax^-/-^Mcl-1^-/-^Vdac2^-/-^* MEFs expressing Cas9 and a whole-genome sgRNA library were cultured with 10 µM ABT-737 or 500 nM A1331852 for 48 h. Surviving cells were expanded for 5 days and sgRNA abundance was quantified by deep-sequencing. Plot shows the abundance of independent sgRNAs in selected versus untreated cells. Values represent the mean of >20 biological replicate samples (2 cell lines x 5-6 independent library transductions subjected to selection with ABT-737 or A1331852 in parallel). Dashed line represents a LOWESS regression curve fit to the data. (**D**) MEFs of the indicated genotypes (*Mcl1*^-/-^ genetic background) were treated with escalating does of the BCL-XL inhibitor A1331852 for 48 h. Cell death was assessed by PI uptake and flow cytometry. Data are mean ± SD of three independent experiments. (**E**) MEFs of the indicated genotypes (*Mcl1*^+/+^ genetic background) were treated with (10 µM MCL-1 inhibitor S63845 and escalating does of A1331852) for 24 h. Cell death was assessed by PI uptake and flow cytometry. Data are mean ± SD of three independent experiments. See also Figure S1.

## RESULTS

### MARCHF5 regulates BAK-dependent apoptosis

To identify genes that regulate BAK-driven apoptosis independently of (or redundantly with) VDAC2, a CRISPR-Cas9 whole genome library screen was performed in *Bax^-/-^ Mcl1^-/-^ Vdac2^-/-^* MEFs. Removing MCL-1 enabled apoptosis to be driven in these cells with the BH3-mimetic compounds ABT-737 or A1331852 (Figure 1A), while the absence of BAX and VDAC2 ensured that apoptosis was mediated by BAK and controlled independently of VDAC2. A genome-wide lentiviral single guide RNA (sgRNA) library^22^ was introduced into *Bax^-/-^ Mcl1^-/-^ Vdac2^-/-^* MEFs. Selective pressure was then applied by culturing the cells with ABT-737 or A1331852 for 48 hours to induce apoptosis. Surviving cells were expanded for 5 days before extracting DNA and amplifying sgRNA sequences (Figure 1B). The most enriched sgRNA sequences were those targeting *Bak* (Figure 1C). This was expected because deleting *Bak* would yield BAK/BAX-deficient cells, which are completely refractory to triggers of intrinsic apoptosis ^31^. Clear enrichment was also observed for multiple sgRNAs targeting two genes involved in ubiquitin signalling: the mitochondrial E3 ubiquitin ligase Membrane Associated RING-CH-type Finger 5 (*Marchf5*) and the ubiquitin receptor ubiquilin-1 (*Ubqln1*) (Figure 1C). The sgRNAs targeting these genes were not enriched in the equivalent screen performed in *Bax^-/-^Mcl1^-/-^Vdac2^+/+^* MEFs ^14^, suggesting that their influence on BAK in these cells becomes most evident when VDAC2 is absent.

To validate the candidates, *Marchf5* or *Ubqln 1* were deleted in *Bax^-/-^ Vdac2^-/-^* MEFs on both an *Mcl1^-/-^* and an *Mcl1^+/+^* genetic background. In both contexts, the *Marchf5* or *Ubqln1* knockout cells were less sensitive to A1331852 compared with *parental* cells transduced with empty vector (Figure 1D,1E, S1H and S1K), confirming the results of the screen. Removing *Marchf5* produced the strongest phenotype in these experiments, so our subsequent studies focused on elucidating the mechanism by which MARCHF5 was influencing BAK.

### Deleting *Marchf5* in combination with *Vdac2* in *Bax^-/-^* MEFs impairs MOMP and cytochrome *c* release

Given that *Bax^-/-^Vdac2^-/-^Marchf5^-/-^* MEFs were highly resistant to treatment with BH3 mimetic drugs in short-term cell death assays, we explored whether this protective effect preserved clonogenic potential. After 5 days treatment with combined BH3 mimetics, there were significantly more colonies formed by *Bax^-/-^Vdac2^-/-^Marchf5^-/-^* MEFs compared with MEFs of other genotypes (*Bax^-/-^* or *Bax^-/-^Vdac2^-/-^* or *Bax^-/-^ Marchf5^-/-^*) (Figure 2A). Mitochondrial outer membrane permeabilization (MOMP) is the point of no return in apoptosis as cells eventually die irrespective of caspase activity following MOMP ^32, 33^. Defects downstream of MOMP (for example APAF-1- or caspase-9-deficiency) typically slow the kinetics of cell death without blocking it entirely ^34,35,36^. Hence, the observation that MARCHF5-deficent cells retain clonogenic potential suggests that the mechanism impairing BAK function likely operates upstream of MOMP.

**Fig. 2.**
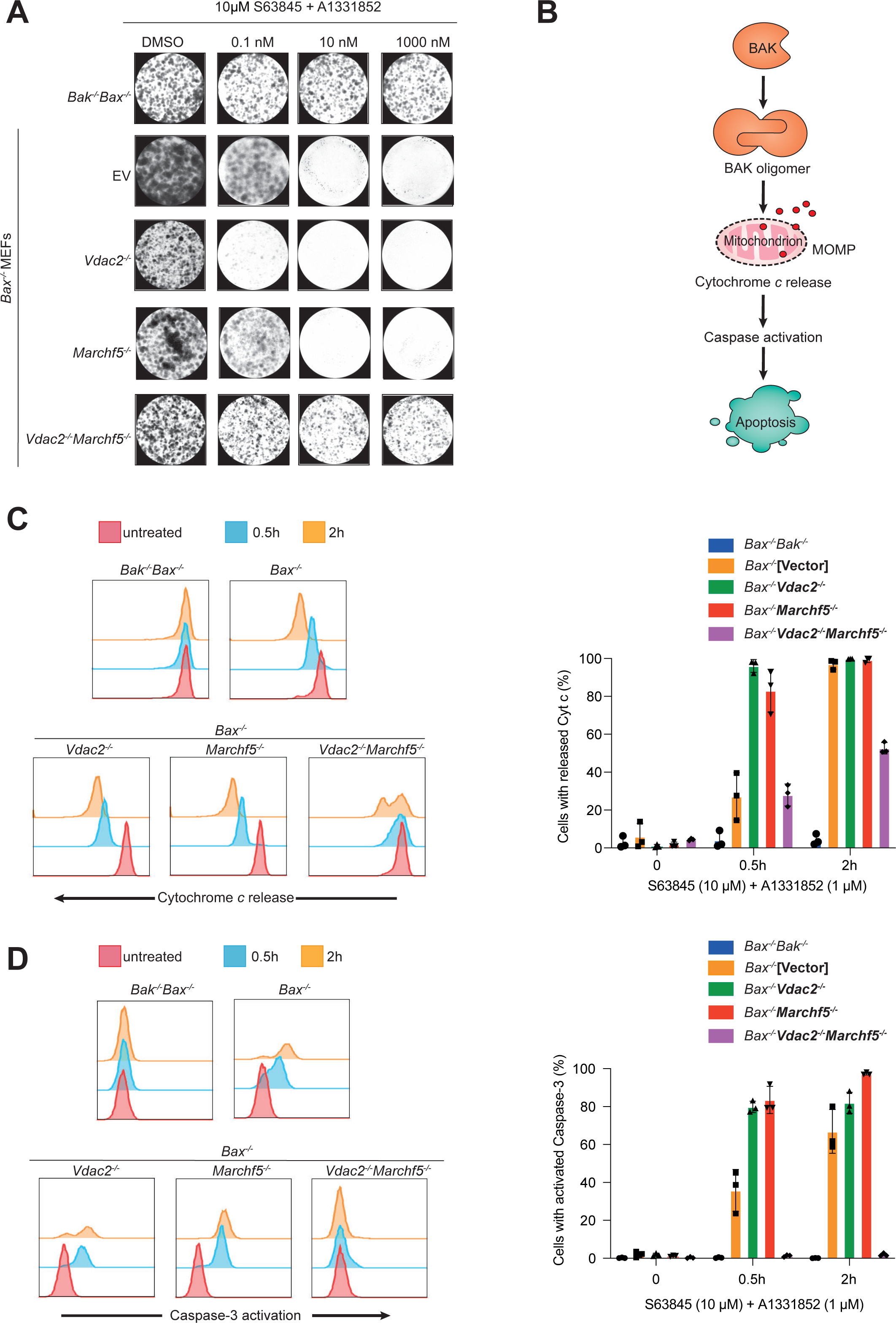
Deleting *Marchf5* in combination with *Vdac2* in *Bax^-/-^* MEFs impairs MOMP and cytochrome *c* release. (**A**) MEFs of the indicated genotypes were cultured continuously with BH3 mimetic drugs (10 µM S63845 and escalating concentration of A1331852) for 5 days. Colonies were stained with crystal violet and photographed. Representative images of three independent experiments are shown. (**B**) Schematic of the intrinsic apoptosis pathway. (**C**) Cells were cultured with BH3 mimetics (10 µM S63845 + 1 µM A1331852) for 0.5 h or 2 h in the presence of 20 µM caspase inhibitor QVD-OPh. Cytochrome *c* release was assessed by immunostaining and flow cytometry. Histograms are representative of three independent experiments and the proportion of cells with low cytochrome *c* collated from three independent experiments, data are mean ± SD. (**D**) Cells were cultured with BH3 mimetics (10 µM S63845 + 1 µM A1331852) for 0.5 h or 2 h. Caspase-3 activation was assessed by immunostaining and flow cytometry. Histograms are representative of three independent experiments with the proportion of cells exhibiting active caspase-3 collated from three independent experiments, data are mean ± SD.

To explore this, we assessed cytochrome *c* release and caspase-3 activation in cells exposed to BH3 mimetic drugs, both of which occur downstream of MOMP (Figure 2B). Following treatment with BH3 mimetics, *Bax^-/-^Vdac2^-/-^ and Bax^-/-^Marchf5^-/-^* MEFs cells exhibited cytochrome *c* release within 2 h. In comparison, fewer *Bax^-/-^Vdac2^-/-^ Marchf5^-/-^* MEFs had released cytochrome *c* at the 2 h timepoint (Figure 2C). These data indicate that MARCHF5 loss impairs BAK-driven MOMP induced by BH3 mimetics in MEFs – but only when VDAC2 is absent.

Consistent with the flow cytometry data, when assessed by subcellular fractionation and Western blotting (Figure S2), BH3 mimetics caused rapid and complete cytochrome *c* loss from the mitochondria of *Bax^-/-^Vdac2^-/-^* and *Bax^-/-^Marchf5^-/-^* MEFs, but incomplete and more gradual loss from *Bax^-/-^Vdac2^-/-^Marchf5^-/-^* MEFs. Interestingly, cytochrome *c* only accumulated to appreciable levels in the cytosolic fractions of the *Bax^-/-^Vdac2^-/-^* and *Bax^-/-^Marchf5^-/-^* MEFs, with only very low levels of cytochrome c observed in the cytosol of *Bax^-/-^Vdac2^-/-^Marchf5^-/-^* MEFs. In all genotypes, a proteasome-dependent (MG-132-inhibited) degradation pathway acted to eliminate cytochrome c once it reached the cytosol ^37^. We hypothesized that the slower kinetics of release, together with this inherent proteasome-turnover pathway, prevented cytochrome c from reaching sufficient levels to activate the caspase cascade in *Bax^-/-^Vdac2^-/-^Marchf5^-/-^* MEFs.

To test this hypothesis, we assessed whether caspase-3 became activated in these cells. Robust caspase-3 activation was observed in *Bax^-/-^Vdac2^-/-^* and *Bax^-/-^Marchf5^-/-^* MEFs exposed to BH3 mimetic drugs, but almost no caspase-3 activation occurred in *Bax^-/-^Vdac2^-/-^Marchf5^-/-^* cells (Figure 2D). These data are consistent with cytochrome c failing to reach a sufficient cytosolic level for caspase activation in the *Bax^-/-^Vdac2^-/-^ Marchf5^-/-^* MEFs. Moreover, we infer that these cells must contain enough intact mitochondria to recover and support clonogenic growth ^38^.

### MARCHF5 deletion alone can be sufficient to regulate BAK-dependent apoptosis

Given our findings in MEFs, we sought to explore if the role of MARCHF5 in regulating BAK was conserved in human cells. We deleted *MARCHF5 and/or VDAC2 in BAX^-/-^* HeLa and KMS-12-PE human multiple myeloma cells using CRISPR/Cas9 gene editing (Figure S3A and S3B). Interestingly, deleting *MARCHF5* alone (i.e independent of *VDAC2* deletion) was sufficient to inhibit BAK-dependent apoptosis in HeLa and KMS-12-PE cells induced by BH3 mimetic drugs in both short-term assays (Figure 3A and S3C) and long-term clonogenic assays (Figure 3B). As in MEFs, in both HeLa and KMS-12-PE cells, *MARCHF5* deletion likewise blocked cell death upstream of MOMP as evidenced by impaired cytochrome *c* release (Figure 3C and S3D) and caspase-3 activation (Figure 3D and S3E).

**Fig. 3.**
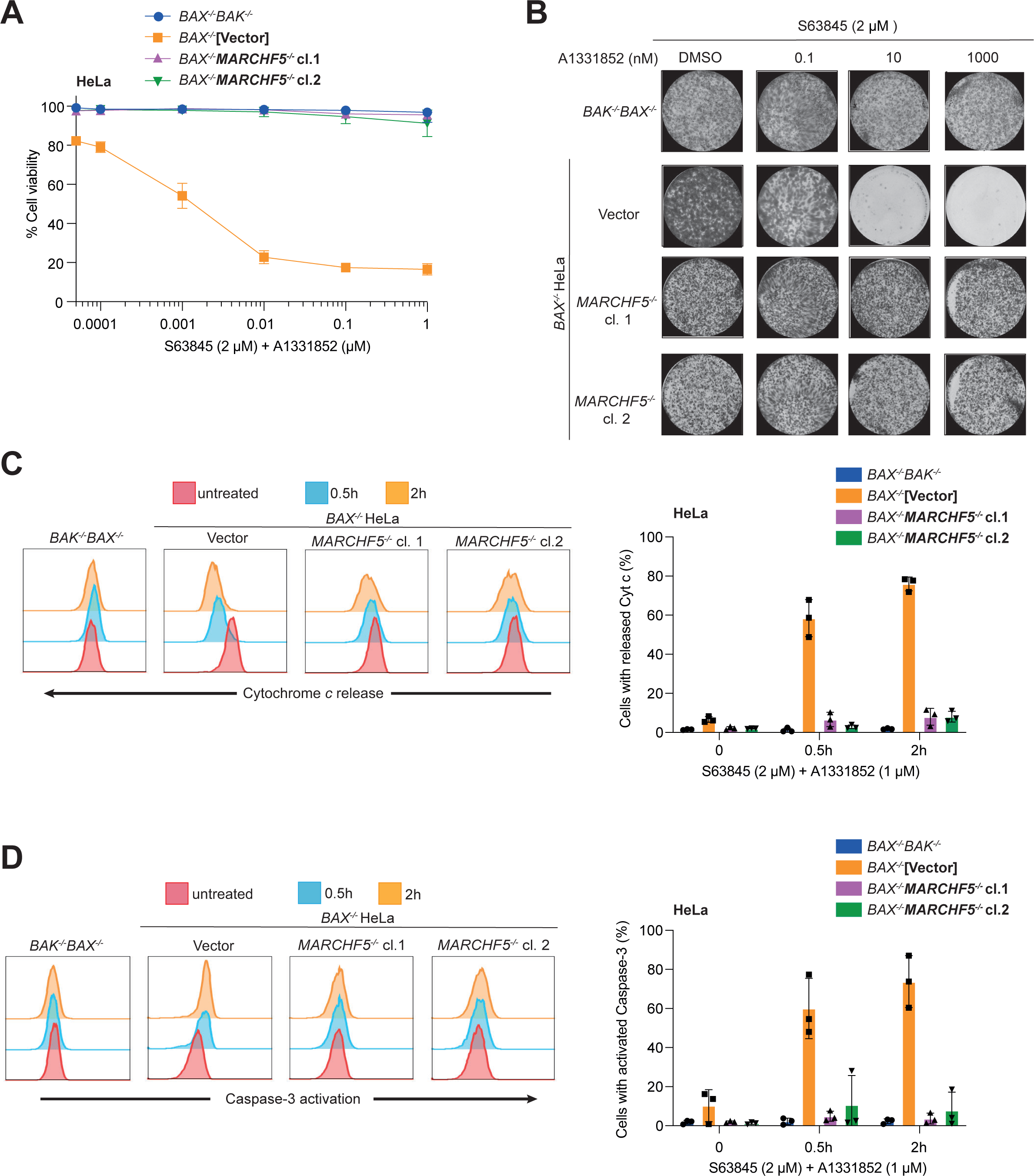
MARCHF5 deletion can be sufficient to regulate BAK-dependent apoptosis. (**A**) HeLa cells of the indicated genotypes were treated with BH3 mimetic drugs (2 µM S63845 and escalating concentration of A1331852) for 24 h. Cell death was assessed by PI uptake and flow cytometry. Data are represented as mean ± SD of three independent experiments. (**B**) Cells were cultured with BH3 mimetic drugs (2 µM S63845 and escalating concentration of A1331852) for 5 days, colonies were stained with crystal violet and photographed. Representative images of three independent experiments. (**C**) Cells were cultured with BH3 mimetic drugs (2 µM S63845 and escalating concentration of A1331852) for 0.5 h or 2 h in the presence of caspase inhibitor QVD-OPh (20 µM). Cytochrome *c* release was assessed by immunostaining and flow cytometry. Representative histograms shown from 3 independent experiments and proportion of cells with low cytochrome *c* collated from three independent experiments, data are mean ± SD. (**D**) Cells were cultured with BH3 mimetic drugs (2 µM S63845 and escalating concentration of A1331852) for 0.5h or 2h. Caspase-3 activation was assessed by immunostaining and flow cytometry. Representative histograms are from 3 independent experiments with proportion of cells exhibiting active caspase-3 collated from three independent experiments, data are mean ± SD. See Figure S3 for results of KMS-12-PE cells.

### *MARCHF5* deletion causes BAK activation and sequestration by pro-survival proteins

Having identified that deleting *Marchf5/MARCHF5* limited BAK-mediated MOMP, we sought to resolve the mechanism. MARCHF5 is an E3 ubiquitin ligase that resides in the mitochondrial outer membrane. It targets a variety of protein substrates for proteasomal turnover by adding K48-linked ubiquitin chains ^17, 39,40,41,42,43,44,44,45,46,47,48^. It can also modulate substrate localisation through K63-linked ubiquitin modifications ^49^. MARCHF5 ubiquitin ligase activity has also been shown to regulate mitochondrial protein import ^50^.

As MARCHF5 is known to target MCL-1 for turnover by the proteasome ^17, 39,40,41,42^, we investigated whether MARCHF5 may effect apoptosis by regulating the abundance and/or mitochondrial localisation of other BCL-2 family proteins. Deleting *MARCHF5* stabilised total MCL-1 and NOXA protein levels in HeLa and KMS-12-PE cells (Figure 4A and Figure S4B), but not in MEFs, which express very low levels of endogenous NOXA (Figure S4A) ^17^. There were no other notable changes in expression level among the BCL-2 family proteins examined (BAK, BCL-XL, BCL-2, PUMA, BAD, BIM) (Figure 4A, Figure S4A and S4B).

**Fig. 4.**
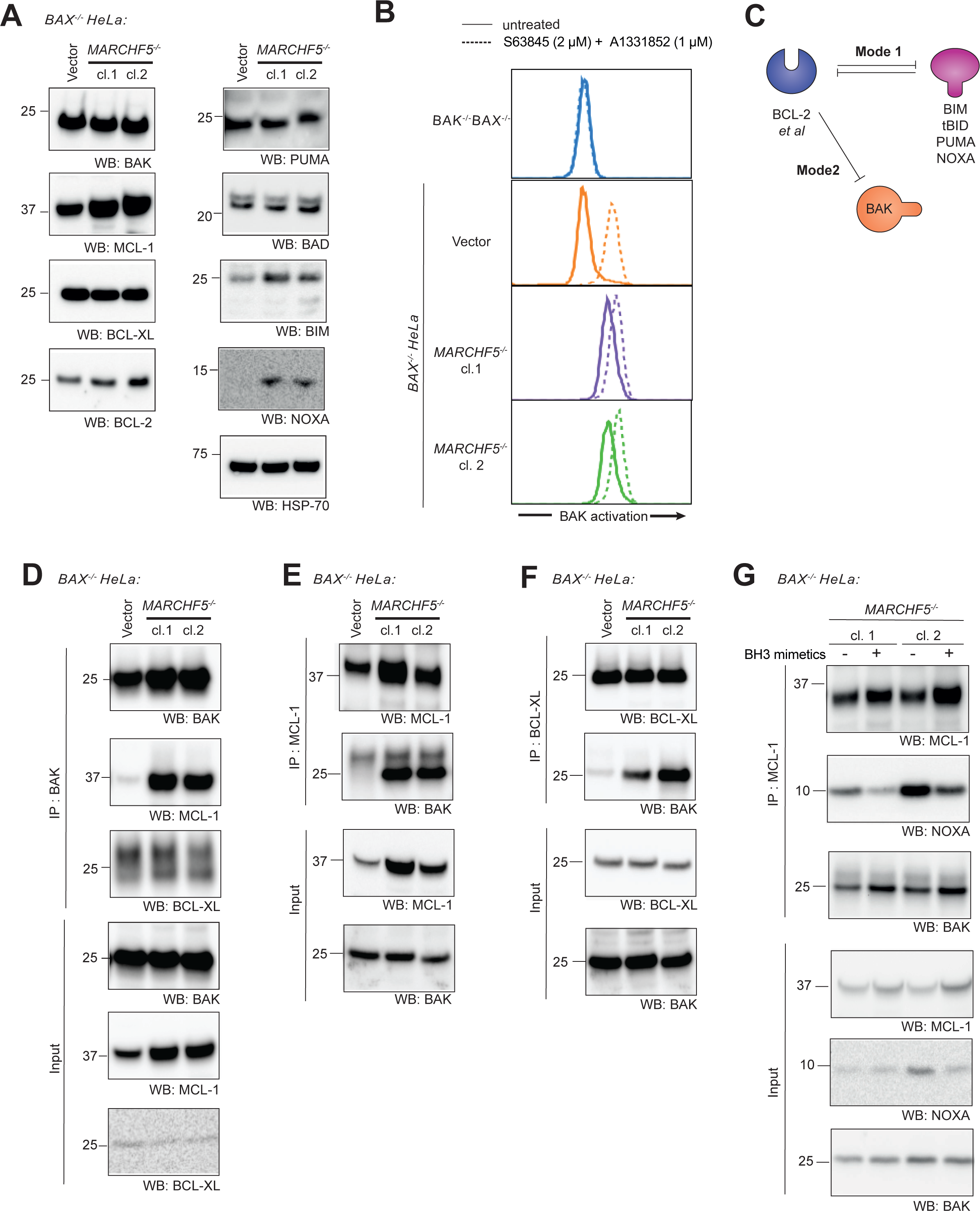
*MARCHF5* deletion causes BAK activation and sequestration by pro-survival proteins. (**A**) BCL-2 family protein abundance was assessed by Western blotting whole cell lysates prepared from HeLa cells of the indicated genotype. Data are representative of two independent experiments. (**B**) Cells were cultured with BH3 mimetic drugs (2 µM S63845 and escalating concentration of A1331852) for 2 h. BAK conformation was assessed by immunostaining and flow cytometry using an antibody that recognises BAK in its active conformation (clone G317-2). Histograms are representative of three independent experiments. (**C**) Schematic representation of inhibitory modes of pro-survival proteins. Cell protection is afforded by sequestering BH3-only proteins (Mode 1) or sequestering activated BAK and BAX (Mode 2). (**D-F**) BAK (**D**), MCL-1 (**E**) or BCL-XL (**F**) were immunoprecipitated from whole cell lysates. Bound proteins were resolved by SDS-PAGE and immunoblotted for BAK, MCL-1 or BCL-XL. Asterisks indicate cross-reactive non-specific bands. Data are representative of three independent experiments. (**G**) MCL-1 was immunoprecipitated from whole cell lysates prepared from untreated cells or cells treated with BH3 mimetic drugs (2 µM S63845 and 1 µM A1331852) for 4 h. Bound proteins were resolved by SDS-PAGE and immunoblotted for BAK, NOXA and MCL-1. Data are representative of three independent experiments. See also Figure S4 for the result of MEFs and KMS-12-PE.

Although MCL-1 levels became elevated following MARCHF5 loss, this was unlikely to be the sole factor impairing BAK because a similar phenotype was observed in *Mcl1-*deficient MEFs (Figure1C and 1D). Nevertheless, we sought to determine whether the elevated levels of MCL-1 served to impair BAK in *MARCHF5^-/-^* HeLa, as it was the most significantly altered BCl_2_ protein in these cells. We assessed the response of *MTCH2*-deficient cells to BH3 mimetics because MTCH2 is also necessary for MARCHF5 to degrade MCL-1 ^17^. Deleting *MTCH2* in *BAX^-/-^* HeLa stabilised MCL-1 to similar levels as *MARCHF5* deletion (Figure S4C). However, in contrast to MARCHF5-deficienct cells, MTCH2-deficiency did not confer resistance to BH3-mimetic drugs (Figure S4D), indicating that MCL-1 stabilisation alone was not responsible for the protection afforded by MARCHF5 loss.

In order to drive apoptosis, BAK undergoes significant conformational rearrangement ^51,52,53,54,55,56,57,58,59,60,61,62^. Given that the apoptosis deficit in MARCHF5-deficient cells is upstream of MOMP, we hypothesized that BAK confirmation change may be perturbed in the absence of MARCHF5. BAK activation was assessed at steady-state and following exposure to BH3 mimetic drugs using intracellular flow cytometry. As expected, very little activated BAK could be observed in untreated MARCHF5 wild-type HeLa (*BAX^-/-^*), MEFs (*Bax^-/-^Vdac2^-/-^*) and KMS-12-PE cells (*BAX^-/-^*), and the amount of active BAK increased following exposure to BH3 mimetics (Figure 4B, S4E and S4F). Surprisingly, however, in MARCHF*5*-deficient cells, BAK constitutively adopted an active conformation even in the absence of BH3 mimetics. When exposed to BH3 mimetic drugs, BAK underwent little if any further activation (Figure 4B, S4E and S4F), suggesting that most of BAK in untreated MARCHF5-deficient cells was already in an active conformation.

Given that BAK failed to drive apoptosis in MARCHF5-deficient cells despite adopting a conformation normally associated with its active form, we hypothesized that it may be sequestered by pro-survival BCL-2 proteins and so unable to execute its killer activity. In BAK-dependent apoptosis, the apoptotic threshold is set by interactions between BCL-2 family proteins. Pro-survival proteins prevent cell death by either interacting with BH3-only proteins (Mode 1) or binding to activated BAK/BAX (Mode 2) (Figure 4C) ^63^. Mode 2 interactions are proposed to be less easily disrupted compared to Mode 1 interactions ^63, 64^ and may confer resistance to anti-cancer treatments ^63, 65,66,67^. It is possible that loss of MARCHF5 impacts on the interactions between BCL-2 family members, which may render the cells less responsive to BH3 mimetic drugs. Hence, we evaluated the interactions between BAK and the pro-survival proteins MCL-1 or BCL-XL by co-immunoprecipitation. Using a conformation-specific antibody to pull down the active from of BAK (clone G317-2) ^55^, greater amounts of both MCL-1 and BCL-XL were co-precipitated from MARCHF5-deficient HeLa than from parental vector control cells (Figure 4D). Likewise, when either MCL-1 or BCL-XL were immunoprecipitated, more BAK co-precipitated from the cells lacking MARCHF5 (Figure 4E and 4F). Similar profiles were also observed in MARCHF5-deficient MEFs (Figure S4G-I) and KMS-12-PE (Figure S4J and S4K). Treatment of MARCHF5-deficient HeLa cells with BH3 mimetics disrupted the Mode 1 interaction between MCL-1 and NOXA, but the Mode 2 MCL-1-BAK protein complexes were retained (Figure 4G). Together, these data suggest that in the absence of MARCHF5, BAK changes conformation and engages in Mode 2 complexes with pro-survival proteins that are not readily be disrupted by BH3 mimetics. Consequently, BAK becomes restricted in its ability to mediate apoptosis.

We noted that MARCHF5-deficient HeLa cells have elevated levels of MCL-1, and that this change could contribute to the enhanced co-precipitation observed with BAK. To control for this change we also performed the co-precipitations in MTCH2-deficient HeLa cells, which have comparably elevated levels of MCL-1 (Figure S4C). In these experiments, we continued to observe significantly more BAK co-precipitating with MCL-1 in *MARCHF5^-/-^* than in wild-type or *MTCH2^-/-^* cells (Figure S4L), indicating that the elevated MCL-1 levels alone are not sufficient to provoke the BAK:MCL-1 Mode 2 interaction.

### MARCHF5 regulates BAK apoptotic activity dependent on its E3 ubiquitin ligase activity

Given that MARCHF5 is an E3 ubiquitin ligase, we next sought to investigate whether its enzymatic activity was key to regulating BAK. We began by engineering HeLa cells expressing MYC-tagged MARCHF5, either in its wild-type active form or a mutant form that lacks ubiquitin ligase activity (MARCHF5^ΔCys^) ^43, 46^. Ectopic expression was confirmed by Western blotting for the Myc tag (Figure 5A). Whilst re-expressing wild-type MARCHF5 rescued the apoptotic response of MARCHF5-deficient cells, those expressing MARCHF5^ΔCys^ remained highly resistant to BH3 mimetic drugs (Figure 5B).

**Fig. 5.**
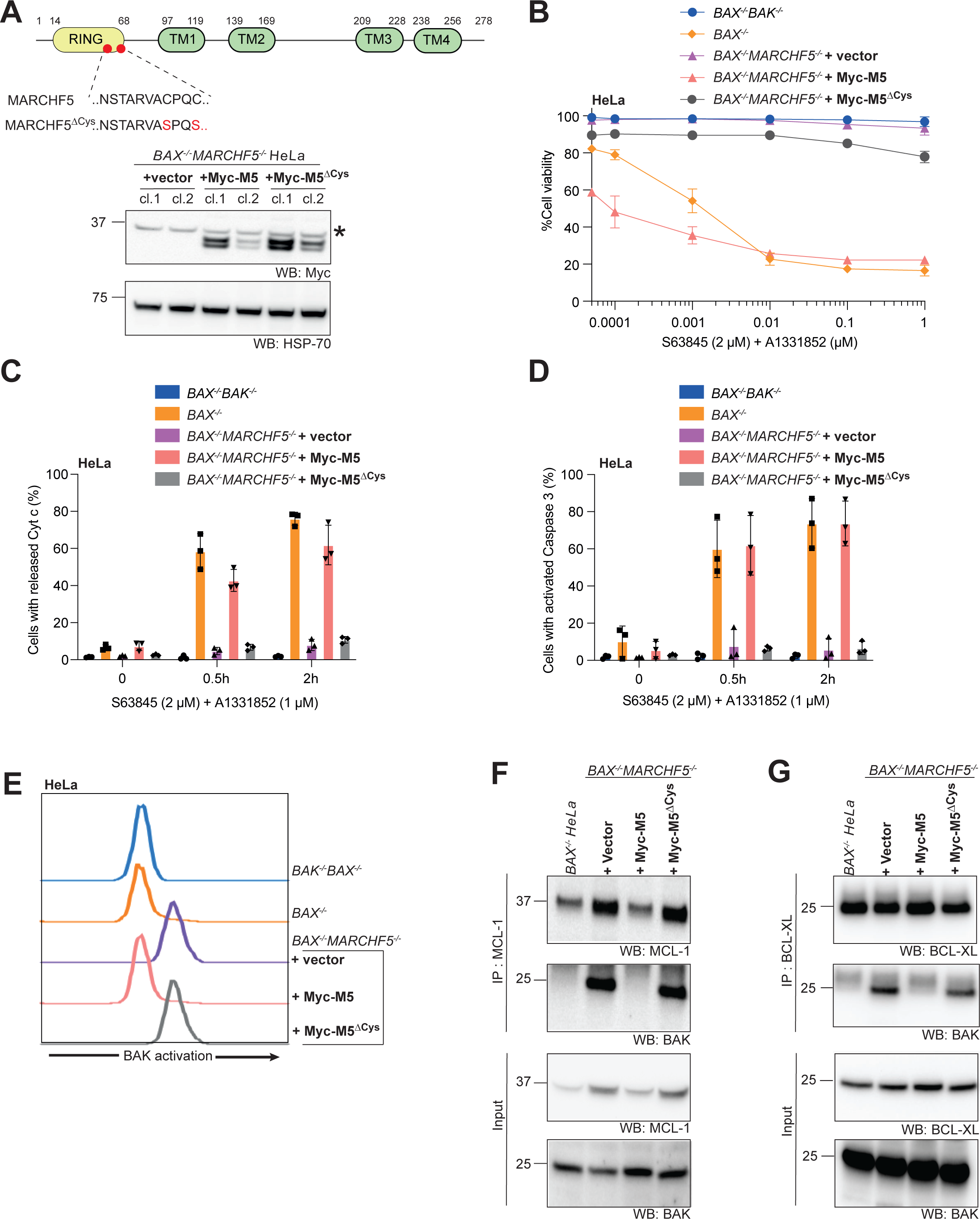
MARCHF5 regulates BAK dependent on its E3 ubiquitin ligase activity. (**A**) MYC-tagged wild-type MARCHF5 or catalytically inactive mutant MARCFH5^ΔCys^ (C65S, C68S) were expressed in *BAX^-/-^MARCHF5^-/-^* HeLa cells. Exogenous MARCHF5 protein expression was confirmed by immunoblotting for the Myc-tag. The asterisk indicates the position of the non-specific bands. Data are representative of three independent experiments. (**B**) The indicated cells were cultured with BH3 mimetic drugs (2 µM S63845 and escalating concentration of A1331852) for 24 h. Cell death was assessed by PI uptake and flow cytometry. Data are mean ± SD of three independent experiments. (**C**) Cells were cultured with BH3 mimetic drugs (2 µM S63845 with 1 µM A1331852) for 0.5 h or 2 h in the presence of caspase inhibitor QVD-OPh (20 µM). Cytochrome *c* release was assessed by immunostaining and flow cytometry. Representative histograms shown from 3 independent experiments and the proportion of cells with low cytochrome c collated from three independent experiments. Data are presented as mean ± SD. (**D**) Cells were cultured with BH3 mimetic drugs (2 µM S63845 and 1 µM A1331852) for 0.5 h or 2 h. Caspase-3 activation was assessed by immunostaining and flow cytometry. Representative histograms are from 3 independent experiments with the proportion of cells exhibiting active caspase-3 collated from three independent experiments. Data are presented as mean ± SD. (**E**) BAK conformation was assessed in the indicated cells by immunostaining and flow cytometry using a BAK conformation-specific antibody (clone G317-2). Histograms are representative of three independent experiments. (**F-G**) MCL-1 (**F**) or BCL-XL (**G**) were immunoprecipitated from whole cell lysates prepared from the indicated cells. Bound proteins were resolved by SDS-PAGE and immunoblotted for BAK, MCL-1 or BCL-XL. Data are representative of three independent experiments.

**Fig. 6.**
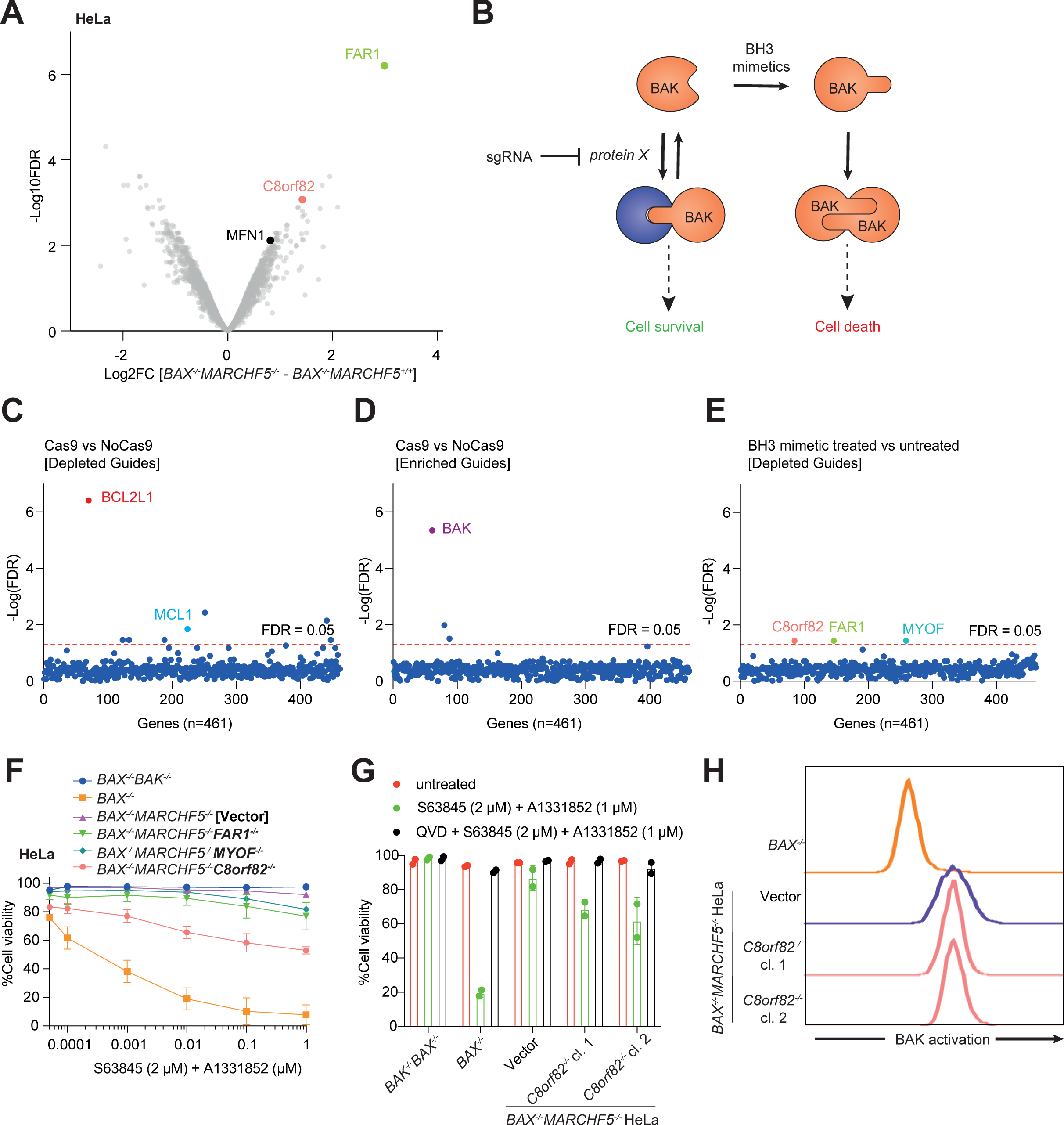
Identification of MARCHF5 substrates that contribute to BAK activation. (A) Volcano plot depicting differences in protein abundance between *BAX^-/-^ MARCHF5^+/+^* and *BAX^-/-^MARCHF5*^-/-^ HeLa cells. See Table S3 for additional details. (B) Schematic of the rationale for the drop-out CRISPR/Cas9 library screen. If BAK is driven into inhibitory Mode 2 complexes by a substrate protein that accumulates upon *MARCHF5* deletion, then deleting the substrate should restore BAK-mediated apoptosis in *MARCHF5* knockout cells. As such, sgRNA targeting the substrate should be depleted following BH3 mimetic treatment. (**C - D**) *BAX^-/-^MARCHF5^-/-^* HeLa cells with or without Cas9 expression were transduced with a custom lentiviral sgRNA library (see Table S5 for additional details). Transduced cells cultured for 5 days. sgRNA were amplified from surviving cells and quantified by sequencing. The significance (FDR-corrected) of gene depletion (**C**) or enrichment (**D**) was calculated using the ranked order of the Log2FC for individual sgRNA targeting each gene ^30^. See Table S6A and S6B for additional detail. (**E**) *BAX^-/-^MARCHF5^-/-^* HeLa cells with Cas9 expression from (**C**) were treated with DMSO or BH3 mimetic drugs (2 µM S63845 and 1 µM A1331852) for 48 h. sgRNA were amplified from surviving cells and quantified by sequencing. Gene-level significance values were calculated as above. See Table S6C for additional detail. (**F**) The cells were cultured with BH3 mimetic drugs (2 µM S63845 and escalating concentration of A1331852) for 24 h. Cell death was assessed by PI uptake and flow cytometry. Data are presented as mean ± SD of three independent experiments. (**G**) *BAX^-/-^MARCHF5^-/-^C8ORF82^-/-^* and control HeLa cells were cultured with BH3 mimetic drugs (2 µM S63845 and 1 µM A1331852) in the presence or absence of caspase inhibitor (QVD-OPh, 50 µM) for 24 h. Cell death was assessed by PI uptake and flow cytometry. Data are presented as mean ± SD of two independent experiments. (**H**) BAK conformation was assessed by immunostaining and flow cytometry in *BAX^-/-^MARCHF5^-/-^ C8ORF82^-/-^* and control HeLa cells using a BAK conformation-specific antibody (clone G317-2). Histograms are representative of two independent experiments.

Consistent with this observation, only the active form of MARCHF5 could restore the ability of BH3 mimetics to cause cytochrome *c* release (Figure 5C) and activate caspase-3 (Figure 5D). Likewise, the propensity for BAK to change conformation (Figure 5E) and engage in Mode 2 interactions with MCL-1 (Figure 5F) and BCL-XL (Figure 5G) were all reversed by the expression of wild-type MARCHF5, but not MARCHF5^ΔCys^. This was also observed in MEFs and KMS-12-PE cell lines (Figure S5). These data suggest that MARCHF5 regulates BAK conformation change and BAK-driven apoptosis via its E3 ubiquitin ligase activity.

### Identification of MARCHF5 substrates that contribute to BAK activation

Our data shows that MARCHF5 ubiquitin ligase activity regulates BAK conformation and function, but we did not observe any changes in BAK stability following MARCHF5 loss (Figure 4A and S4A), suggesting that BAK is unlikely to be a direct substrate. Instead, we hypothesise that MARCHF5 regulates BAK indirectly. MARCHF5 targets many substrates for protein degradation through the addition of K48-linked ubiquitin ^17, 39, 43,44,45,46,47,48, 68^. Upon *MARCHF5* deletion, some of these substrate proteins may accumulate and thereby influence BAK activation and its interaction with pro-survival proteins.

To identify proteins that may be responsible for driving BAK into inhibitory Mode 2 complexes in MARCHF5-deficient cells, we used two complimentary approaches. Firstly, we compared the proteome of MARCHF5 wild-type and MARCHF5-deficient cells by mass spectrometry, using both HeLa and MEFs. We identified 32 proteins that were significantly elevated in *BAX^-/-^MARCHF5^-/-^* HeLa (Log2FC > 1; FDR < 0.05) (Figure 6A, S6A and Table S3). The known MARCHF5 substrate MFN1 was significantly enriched in this dataset using a slightly less stringent Log2FC cut-off (Log2FC = 0.81; FDR = 0.008) ^43,44,45^ (Figure 6A and Table S3). Sample preparation was slightly more variable for the MEF proteomics dataset, resulting in fewer proteins reaching a statistically significant threshold, but three proteins stood out as being elevated in *Vdac2^-/-^Marchf5^-/-^* relative to *Vdac2^-/-^Marchf5^+/+^* MEFs (Log2FC > 1; FDR = 0.05) (Figure S6B, S6C and Table S4). Interestingly, one of those proteins was fatty acyl CoA reductase 1 (FAR1), which was also the most significantly elevated protein in *MARCHF5^-/-^* HeLa (Log2FC = 3.0; FDR = 6.3 x 10^-7^) (Figure 6A, S6A and Table S3).

To explore whether the any of these elevated proteins was responsible for the impaired BAK activity in MARCHF5-deficient cells, we generated a custom sgRNA library to perform CRISPR/Cas9 loss-of-function dropout screens. In addition to the proteins found to be elevated in our proteomic analyses, we also included sgRNA targeting previously reported MARCHF5 substrates, peroxisomal proteins and other known components of the MOM. The final library comprised 3325 sgRNAs targeting 461 genes (6 or 7 sgRNAs for each gene) and 150 non-targeting control sgRNAs (Table S5).

The library was introduced by lentiviral infection into *BAX^-/-^MARCHF5^-/-^* HeLa that either did or did not express Cas9 endonuclease. To identify sgRNA targeting genes that influenced the steady-state growth and survival of *BAX^-/-^MARCHF5^-/-^* HeLa, we compared changes in sgRNA abundance between Cas9^+ve^ cells to Cas9^-ve^ cells. We observed a significant depletion of sgRNA targeting *BCl_2_L1/BCLXL* and *MCL1* in cells expressing Cas9 (Figure 6C and S6D), indicating that these proteins were necessary to maintain survival of *BAX^-/-^MARCHF5^-/-^* cells, likely due to their capacity to form Mode-2 interactions that restrain BAK. Conversely, sgRNAs targeting BAK were enriched in this comparison (Figure 6D and S6D), indicating that removing BAK conferred a survival advantage.

Next, we sought to identify sgRNA that would reverse the impaired response to BH3 mimetics caused by MARCHF5-deficiency in *BAX^-/-^* HeLa. The *BAX^-/-^MARCHF5^-/-^* cells expressing Cas9 and the sgRNA library were subjected to selection with BH3-mimetics (2 µM S63845 +1 µM A1331852) for 48 h. sgRNAs targeting 3 genes were significantly depleted following selection (FDR < 0.05): *FAR1*, *C8orf82* and *MYOF* (Figure 6E and S6E). These data suggest that depletion of these genes may re-sensitize *BAX^-/-^MARCHF5^-/-^* cells to apoptosis, marking them as potential BAK-regulating substrates of MARCHF5.

FAR1 and C8orf82 were of particular interest given that these were among the most upregulated proteins following MARCHF5 deletion in our proteomic analyses (Figure 6A and S6A-C). To validate whether any of these proteins could regulate BAK apoptotic activity, each was deleted in *BAX^-/-^MARCHF5^-/-^* HeLa by CRISPR/Cas9 gene editing using two independent sgRNA (Figure S6F-H). Deleting *FAR1* or *MYOF* alone did not markedly affect the response of *BAX^-/-^MARCHF5^-/-^* cells to BH3 mimetics (Figure 6F). Intriguingly, deleting *C8orf82* partially restored caspase-mediated apoptosis in *BAX^-/-^MARCHF5^-/-^* MEFs in response to BH3 mimetics (Figure 6F and 6G). However, the loss of C8orf82 did not reverse the constitutive activation of BAK (Figure 6H), suggesting that C8orf82 was not responsible for this aspect of the MARCHF5-deficient phenotype and might instead influence BAK-mediated apoptosis through a different mechanism.

### MARCHF5 controls BAK independently of BH3-only proteins

Our proteomics and genetic screening did not reveal a strong candidate for the MARCHF5 substrate regulating BAK. However, these approaches were predicated on the idea that this substrate would normally be degraded by MARCHF5 and accumulate in its absence. In addition to targeting substrates for proteasomal degradation, MARCHF5 can also mark substrates with forms of ubiquitin that impact localization or function rather than protein stability. If MARCHF5 was acting in this way, then other BCL-2 proteins known to regulate BAK may still represent potential substrates, even though we did not observe notable changes in their abundance (Figure 4A, S4A and S4B). Engagement with BH3-only proteins, such as BID, is typically regarded as necessary to initiate BAK activating conformation change. Therefore, we sought to test whether BH3-only proteins were critical for driving BAK into Mode 2 inhibitory complexes following MARCHF5 deletion. For these experiments we utilised HCT-116 cells that have been engineered to lack all known BCL-2 proteins, including the eight BH3-only family members ^16^. BAK and MCL-1 were re-introduced to these cells with retroviral expression vectors (Figure 7B), and *MARCHF5* was subsequently removed by CRISPR/Cas9 gene editing (Figure S7). In vector control cells, where MARCHF5 expression is maintained, little interaction was observed at steady state between MCL-1 and BAK (Figure 7C). Much greater levels of BAK:MCL-1 Mode 2 interaction were observed in two independent *MARCHF5^-/-^* clones (Figure 7C). These data suggest that BH3-only proteins are not required to drive BAK to the active from that engages in Mode 2 interactions when MARCHF5 is absent.

**Fig. 7.**
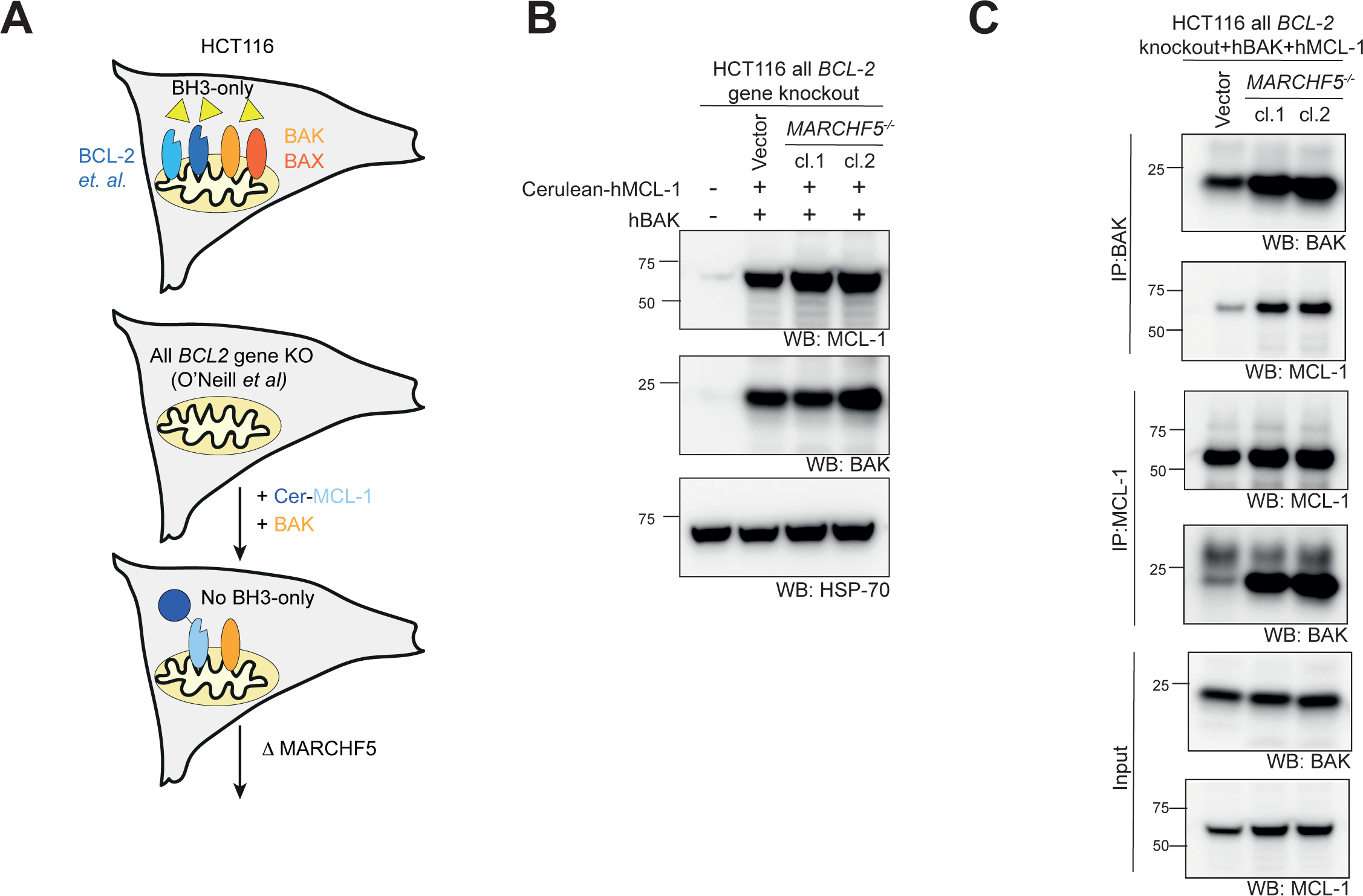
MARCHF5 controls BAK conformation change independently of BH3-only proteins. (**A and B**) hBAK and Cerluean-MCL-1 were stably expressed in HCT-116 cells lacking all BCL-2 family genes ^16^. Protein expression was confirmed by immunoblotting. Two independent *MARCHF5^-/-^* clones were generated (see Figure S7 for indel confirmation). Data are representative of two independent experiments. (**C**) BAK or MCL-1 were immunoprecipitated from whole cell lysates prepared from *MARCHF5^+/+^* or *MARCHF5^-/-^ BCL-2 gene all knockout* HCT-116 cells re-expressing cerulean-hMCL-1 and hBAK. Bound proteins were resolved by SDS-PAGE and immunoblotted for BAK, MCL-1 or BCL-XL. Data are representative of two independent experiments.

## DISCUSSION

MARCHF5 has been linked to mitochondrial apoptosis because it can ubiquitinate the pro-survival protein MCL-1 ^17, 39,40,41,42^, and several proteins that regulate mitochondrial morphology ^48, 69, 70 43,44,45,46,47,48,49^. *MARCHF5*-deficient cells exhibit a hyper-fused mitochondrial network and diminished sensitivity to death stimuli in some cellular contexts ^48, 69, 70^. In other cases, the loss of MARCHF5 can promote apoptosis even though MCL-1 protein levels are elevated ^17, 39, 42^. Here, we have identified that loss of MARCHF5 renders cells resistant to BAK-driven apoptosis (Figure 1D, 1E, 2A, 3A, 3B and S3C). We describe a molecular basis for this observation whereby BAK adopts an active conformation and forms inhibitory Mode 2 complexes in the absence of MARCHF5 ubiquitin ligase activity.

Intriguingly, whilst it was necessary to co-delete *Vdac2* in MEFs to reveal the BAK phenotype caused by MARCHF5-deficiency (Figure 1D, 1E and 2A), deleting *MARCHF5* alone was sufficient to render *BAX^-/-^* HeLa and KMS-12-PE cells resistant to BH3 mimetic drugs (Figure 3A, 3B and S3C). This difference may relate to disparities in how mouse and human BAK interact with VDAC2, as noted from the divergent impact of small molecules on these interactions ^71^. However, other factors may also be involved, and follow-up studies in additional contexts would help to determine whether species-differences in BAK:VDAC2 interaction contribute to the observed phenotypes.

An interesting aspect of the phenotype that results from MARCHF5 loss is that BAK appears to be driven into its active conformation, but paradoxically this is associated with diminished sensitivity to apoptotic stimuli. In most cases BAK activation and conformation change would drive cell death through the formation of BAK dimers and higher-order oligomers that damage the mitochondrial outer membrane ^52, 56, 58, 72^. In cells lacking MARCHF5, conformationally altered BAK preferentially engages with pro-survival proteins in inhibitory Mode 2 complexes instead of forming homodimers (Figure 4D-F). Whether this results from changes at the mitochondria that actively suppress BAK-dimerization or ones that encourage Mode 2 interactions is not clear, but elucidating the underlying mechanism could reveal novel ways to modulate apoptosis signalling.

Our studies focused on BAK-driven apoptosis by using cells that did not express any BAX protein. In this context, BAK function was markedly reduced upon loss of MARCHF5. This is in distinct contrast to studies in cells expressing both BAX and BAK, where MARCHF5 deletion resulted in greater sensitivity to apoptosis triggers ^17, 39, 41^. One hypothesis to reconcile these findings is that deleting MARCHF5 could also promote BAX-activating conformation change, but without driving BAX into inhibitory Mode 2 interactions. That could explain for the observed differences but requires further exploration.

The E3 ubiquitin ligase activity of MARCHF5 is specifically what prevents BAK from becoming locked into inactive Mode 2 complexes with pro-survival proteins (Figure 5). This suggests that the ubiquitination of one or more MARCHF5 substrate proteins controls the observed phenotype. Our proteomics and genetic screens failed to highlight a compelling candidate, and it remains uncertain whether ubiquitination by MARCHF5 influences the stability, localization or function of the relevant substrate(s). The substrate may directly influence BAK and/or its pro-survival counterparts, but could equally influence these proteins indirectly, perhaps by altering the mitochondrial lipid environment in which they reside. Mitochondrial lipids can actively participate in the regulation of BAX/BAK-driven apoptosis ^73,74,75^, and MARCHF5 is known to influence lipid and fatty acid metabolism as well as mitochondrial lipid modifications ^76, 77^. There could also be interplay in this regard with the dominant influence of VDAC2 over the phenotype in certain contexts (Figure 1E), given the propensity for VDAC2 loss to disrupt peroxisomes, which are themselves major sites of lipid metabolism ^78^.

It is notable that BAK was driven to its active form in cells lacking MARCHF5 independently of its canonical activators - the BH3-only proteins (Figure 7C). This finding aligns with observations that BAK and BAX can be driven to their active conformation solely through interaction with mitochondrial lipids ^16^. Changes in the nature or composition of mitochondrial lipids following MARCHF5 deletion ^76, 77^ could conceivably impact the propensity for BAK to become activated as well as influence whether activated BAK molecules form homodimers or engage with pro-survival proteins.

Altogether, these findings uncover a hitherto unappreciated mechanism for BAK to transition to its activated conformation, which only becomes apparent upon the removal of MARCHF5 E3 ligase activity. Our studies raise several interesting questions about the mechanisms underlying this process, including the identity of the MARCHF5 substrates responsible and why BAK molecules activated in this way preferentially engage with their pro-survival counterparts in lieu of forming homodimers that would drive cell death. Answering these questions may provide important new insights into the molecular control of apoptosis signalling.

## Acknowledgements

We would like to thank Ruth Kluck, Peter Czabotar, Daniel Gray for helpful discussions and suggestions; Melissa Shi for technical assistance; Stephen Wilcox for DNA sequencing; Simon Cobbold for assistance with proteomic analysis; Prof Xu Luo for providing all *BCL-2* genes knockout HCT116 cells; Christine White for technical support. We thank the Bio21 Mass Spectrometry and Proteomics Facility (MMSPF) for the provision of instrumentation, training, and technical support, and the Mito Foundation for the provision of instrumentation through the large equipment grant support scheme. This work was supported by scholarship fellowship and grants from the Australian National Health and Medical Research Council (NHMRC) to DAS (#1140851, #1140906); Melbourne University (MIRS and MIFRS scholarships to SAH and TMD); the Walter and Eliza Hall Institute of Medical Research (to SAH); the Bodhi Foundation (to GD). Research was supported by an NHMRC Independent Research Institutes Infrastructure Support Scheme grant (361646) and Victorian State Government Operational Infrastructure Support grant.

## Author contributions

DCSH, MvD, and GD devised the study and interpreted data. SAH and HSC planned, performed experiments and interpreted data. BR and DAS performed the proteomics and interpreted data. TMD and IKLT generated MARCHF5 expression constructs; SAH and MvD collated and analysed data. SAH, MvD, GD wrote the manuscript. All authors contributed to manuscript review.

## Supplementary Figure Legends

**Figure S1. Genome-wide CRISPR-Cas9 library screen identifies MARCHF5 and UBQLN1 as regulators of BAK-mediated apoptosis.**

(**A**) Top 10 genes with enriched independent sgRNA in ABT-737 resistant cells. sgRNA were ranked by their Log_2_ fold change (residuals to the LOWESS curve in Fig. 1C). Gene-level significance values were determined from the rank order of sgRNA using an iterative hypergeometric test. (**B-E**) *Marchf5* was deleted from *Bax^-/-^Mcl-1^-/-^Vdac2^-/-^*MEFs (**B**), *Vdac2* was deleted from *Bax^-/-^* MEFs (**C**) and *Marchf5* was deleted from either *Bax^-/-^* (**D**) or *Bax^-/-^Vdac2^-/-^* MEFs (**E**) by CRISPR/Cas9 gene editing using one or two independent sgRNAs derived from the whole genome sgRNA library. DNA sequencing was performed on independent single-cell derived clones to confirm indel generation at sgRNA targeting sites, as illustrated. In some clones, more than two alleles are identified reflecting aneuploidy at the targeted loci. sgRNA target sequences are highlighted in red, and the size of indels is indicated at the end of each sequence, ‘-’ for deletion and ‘+’ for insertion. Clones were selected only if the indels on all alleles detected produced frameshift loss-of-function mutations. (**F, G** and **H**) *Ubqln1* was deleted from *Bax^-/-^Mcl-1^-/-^Vdac2^-/-^*MEFs by CRISPR/Cas9 gene editing using two independent sgRNAs derived from the whole genome sgRNA library. Successful *Ubqln1* mutation in single cell-derived clones was confirmed by sequencing (**F**) and the loss of protein expression (**G**). The asterisk indicates the position of the non-specific bands. Blots are representative of two independent experiments. MEFs of the indicated genotypes (*Mcl1*^-/-^ background) were treated with escalating does of the BCL-XL inhibitor A1331852 for 48 h (**H**). Cell death was assessed by PI uptake and flow cytometry. Data are mean ± SD of three independent experiments. (**I, J** and **K**) *Ubqln1* was deleted from *Bax^-/-^*MEFs (**I**) or *Bax^-/-^Vdac2^-/-^* MEFs (**J**) by CRISPR/Cas9 gene editing using sgRNA derived from the whole genome sgRNA library. Successful *Ubqln1* knock out cells were chosen based on loss of protein expression. The asterisk indicates the position of the non-specific bands. MEFs of the indicated genotypes (*Mcl1*^+/+^ background) were treated with (10 µM MCL-1 inhibitor S63845 and escalating does of A1331852) for 24 h (**K**). Cell death was assessed by PI uptake and flow cytometry. Data are mean ± SD of three independent experiments.

**Figure S2. Deleting *Marchf5* in combination with *Vdac2* in *Bax^-/-^* MEFs impairs cytochrome *c* release.** MEFs of the indicated genotypes were treated with BH3 mimetics (10 µM S63845 + 1 µM A1331852) for 0.5 h or 2 h in the presence of 20 µM caspase inhibitor QVD-OPh. Cells were permeabilized and then separated into cytosol and membrane fractions after treatment. Solubilized proteins from each fraction were run on SDS-PAGE as described. Blots were probed for cytochrome *c*, HSP70 or VDAC1. Data are representative of two independent experiments

**Figure S3 MARCHF5 regulates BAK-dependent apoptosis in HeLa and KMS-12-PE multiple myeloma cells.** (**A-B**) *MARCHF5* was deleted from *BAX^-/-^* HeLa (**A**) and *BAX^-/-^* KMS-12-PE cells (**B**) by CRISPR/Cas9 gene editing using one or two independent sgRNAs. DNA sequencing was performed to identify independent clones with indels causing loss-of-function frameshift mutation at the sgRNA target sites, as illustrated. sgRNA target sequences are highlighted in red, and the size of indels is indicated at the end of each sequence, ‘-’ for deletion and ‘+’ for insertion. (**C**) KMS-12-PE cells of the indicated genotypes were treated with BH3 mimetic drugs (2 µM S63845 with escalating concentration of A1331852) for 24 h. Cell death was assessed by PI uptake and flow cytometry. Data are mean ± SD of three independent experiments. (**D**) KMS-12-PE cells of the indicated genotypes were treated with BH3 mimetic drugs (2 µM S63845 with escalating concentration of A1331852) for 0.5 h or 2 h in the presence of 20 µM Caspase inhibitor QVD-OPh. Cells were analyzed for cytochrome *c* release by flow cytometry. Histograms are representative of three independent experiments and proportion of cells with low cytochrome c collated from three independent experiments, data are mean ± SD. (**E**) KMS-12-PE cells of the indicated genotypes were treated with BH3 mimetic drugs (2 µM S63845 with escalating concentration of A1331852) 0.5 h or 2 h. Cells were analyzed for active caspase-3 by flow cytometry. Histograms are representative of three independent experiments with proportion of cells exhibiting active caspase-3 collated from three independent experiments, data are mean ± SD.

**Figure S4 M*A*RCHF5 deletion drives BAK conformation change to promote its interaction with pro-survival proteins.** (**A**) The protein level of BAK, pro-survival proteins (MCL-1, BCL-XL, BCL-W, BCL-2) and BH3-only proteins (PUMA, BAD, BIM) were analyzed by immunoblotting in MEF cells of the indicated genotypes. The asterisk indicates the position of the non-specific bands. Data are representative of three independent experiments. (**B-C**) The protein level of MCL-1 and NOXA were analyzed by in KMS-12-PE (**B**) and HeLa (**C**) cells of the indicated genotypes. Data are representative of two independent experiments. (**D**) HeLa cells of the indicated genotypes were treated with BH3 mimetic drugs (2 µM S63845 with escalating concentration of A1331852) for 24 h. Cell death was assessed by PI uptake and flow cytometry. Data are presented as mean ± SD of three independent experiments. (**E-F**) MEF (**E**) and KMS-12-PE (**F**) cells of the indicated genotypes were treated with BH3 mimetic drug (10 µM S63845 + 1 µM A1331852 for MEF, 2 µM S63845 + 1 µM A1331852 for KMS-12-PE) for 2 h in the presence of 20 µM Caspase inhibitor QVD-OPh. Cells were then analyzed for BAK conformation change with the conformation-specific antibody G317-2 and assessed by intracellular flow cytometry. Histograms are representative of three independent experiments. (**G-I**) BAK (**G**), MCL-1(**H**) or BCL-XL (I) immunoprecipitations were performed on whole cell lysate prepared from MEF cells. Immunoprecipitates were run on SDS-PAGE and immunoblotted for BAK, MCL-1 or BCL-XL. The asterisk indicates the position of the non-specific bands. Data are representative of three independent experiments. (**J** and **K**) BAK (**J**) or MCL-1(**K**) immunoprecipitations were performed on whole cell lysate prepared from KMS-12-PE cells. Immunoprecipitates were resolved by SDS-PAGE and immunoblotted for BAK or MCL-1. Data are representative of three independent experiments. (**L**) BAK immunoprecipitations were performed on whole cell lysate prepared from HeLa cells. Immunoprecipitates were resolved by SDS-PAGE and immunoblotted for BAK or MCL-1. Data are representative of two independent experiments.

**Figure S5 MARCHF5 regulates BAK through its E3 ubiquitin ligase activity.** (**A** and **B**) MYC-tagged MARCHF5 or catalytically inactive mutant MARCFH5^ΔCys^ (C65S, C68S) were expressed in *Bax^-/-^Marchf5^-/-^* MEFs (two independent clones) and *Bax^-/-^ Vdac2^-/-^Marchf5^-/-^* MEFs (two independent clones) (**A**) or KMS-12-PE *BAX^-/-^ MARCHF5^-/-^* (two independent clones) (**B**). Expression was confirmed by immunoblotting against the Myc-tag. The asterisk indicates the position of the non-specific bands. Data representative of three independent experiments. (**C** and **D**) MEFs (**C**) and KMS-12-PE (**D**) cells of the indicated genotypes were treated with BH3 mimetic drugs (10 µM S63845 with escalating concentration of A1331852 for MEFs and 2 µM S63845 with escalating concentration of A13318522 for KMS-12-PE) for 24 h. Cell death was assessed by PI uptake and flow cytometry. Data are mean ± SD of three independent experiments. (**E** and **F**) MEFs (**E**) and KMS-12-PE cells (**F**) of the indicated genotypes were analyzed for the active conformer of BAK with the conformation-specific antibody G317-2 and assessed by intracellular flow cytometry. Histograms are representative of three independent experiments. (**G** and **H**) BAK(**G**) or MCL-1(**H**) immunoprecipitations were performed on whole cell lysate prepared from MEFs. Immunoprecipitates were resolved by SDS-PAGE and immunoblotted for BAK, MCL-1 or BCL-XL. Data are representative of three independent experiments. (**I** and **J**) MEFs (**I**) and KMS-12-PE cells (**J**) of the indicated genotypes were treated with BH3 mimetic drugs (10 µM S63845 + 1 µM A1331852 for MEFs and 2 µM S63845 + 1 µM A1331852for KMS-12-PE) for 0.5 h or 2 h in the presence of 20 µM Caspase inhibitor QVD-OPh). Cells were analyzed for cytochrome *c* release by flow cytometry. Proportion of cells with low cytochrome *c* collated from three independent experiments and data presented as mean ± SD. (**K** and **L**) MEFs (**K**) and KMS-12-PE cells (**L**) of the indicated genotypes were treated with BH3 mimetic drugs (10 µM S63845 + 1 µM A1331852 for MEFs and 2 µM S63845 + 1 µM A1331852 for KMS-12-PE) for 0.5 h or 2 h. Cells were analyzed for active caspase-3 by flow cytometry. Proportion of cells exhibiting active caspase-3 collated from three independent experiments, data are presented as mean ± SD.

**Figure S6 Identification of MARCHF5 substrates that contribute to BAK conformation change.** (**A**) Heatmap highlighting the relative expression level for proteins that exhibited significant differences in abundance between *MARCHF5^+/+^* and *MARCHF5^-/-^* HeLa cells (FDR ≤ 0.05). Colour scale: fold change relative to the mean peptide intensity in BAX^-/-^ HeLa. (**B**) Volcano plots summarising the magnitude and significance of differences in protein abundance between *Bax^-/-^Vdac2^-/-^Marchf5^+/+^* and *Bax^-/-^Vdac2^-/-^Marchf5^-/-^* MEFs. (**C**) Heatmap highlighting the relative expression level for proteins that exhibited significant differences in abundance between *Marchf5^+/+^* (the mean of *Vdac2^-/-^* and *Bax^-/-^Vdac2^-/-^*) and *Marchf5* knockout (*Bax^-/-^Vdac2^-/-^Marchf5^-/-^*) MEFs (FDR ≤ 0.05). Colour scale: fold change relative to the mean peptide intensity in *wild-type* and *Bax^-/-^* MEFs. (**D**) sgRNAs targeting *MCL-1* or BCl_2_L1 (*BCL-XL)* were depleted while sgRNAs targeting *BAK* were enriched in *BAX^-/-^MARCHF5^-/-^* HeLa with Cas9 expression compared with non-Cas9-expressing cells in the absence of BH3 mimetic drugs. Plot shows the fold change for independent sgRNAs in Cas9 expressing versus non Cas9 expressing cells. Significance values for enrichment or depletion of sgRNA targeting each gene in the library was calculated as described ^30^. See also Table S6 A and B. (**E**) Independent sgRNAs targeting *C8orf82*, *FAR1* or *MYOF* were depleted after treatment with BH3 mimetic drugs compared with non-treated cells. Plot shows the fold change for independent sgRNAs in BH3 mimetic treated versus non treated cells. Significance values for enrichment or depletion of sgRNA targeting each gene in the library was calculated as described ^30^. See also Table S6 C. (**F**) *FAR1* was deleted from *BAX^-/-^MARCHF5^-/-^* HeLa by CRISPR/Cas9 gene editing using two independent sgRNAs. Successful *FAR1* knock out cells were chosen based on loss of protein expression. The asterisk indicates non-specific bands. (**G** and **H**) *MYOF* (**G**) or *C8orf82* (**H**) were deleted from *BAX^-/-^MARCHF5^-/-^* HeLa by CRISPR/Cas9 gene editing using two independent sgRNAs. DNA sequencing was performed to identify independent clones indel with indels causing loss-of-function frameshift mutations. The sgRNAs targeting sequence is highlighted in red, and the size of indels is indicated at the end of each sequence, ‘-’ for deletion and ‘+’ for insertion.

**Figure S7 Deletion of MARCHF5 from all *BCL-2* gene knockout HCT116 cells that express Cerulean-hMCL-1 and hBAK.** *MARCHF5* was deleted from all *BCL-2* gene knockout HCT-116 cells that expressed Cerulean-hMCL-1 and hBAK by CRISPR/Cas9 gene editing. DNA sequencing was performed on independent clones to confirm loss-of-function indel generation at sgRNA targeting sites as shown. The sgRNAs targeting sequence is highlighted in red, and the size of indels is indicated at the end of each sequence, ‘-’ for deletion and ‘+’ for insertion.

